# High-Dimensional Immunophenotyping of Murine T-cell and B-cell Subsets

**DOI:** 10.1101/2020.05.07.082214

**Authors:** Kyle T. Mincham, Jacob D. Young, Deborah H. Strickland

## Abstract

**Purpose and appropriate sample types:** This 19-parameter, 18-colour flow cytometry panel was designed and optimised to enable the comprehensive and simultaneous immunophenotyping of distinct T-cell and B-cell subsets within murine lymphoid tissues (Table 1). Cellular populations identified by employing this OMIP include 4 major subsets of B-cells (memory, activated, plasma cells and plasmablasts) and 7 major subsets of CD4^+^ T-cells (naïve, central memory, effector memory, helper, regulatory, follicular helper and follicular regulatory). Staining was performed on freshly isolated splenocytes from 21-day-old neonatal BALB/c mice, however due to the omission of mouse strain-specific markers, this OMIP can be implemented across a range of murine models where in-depth immunophenotyping of the diverse repertoire of T-cell and B-cell populations localised within lymphoid tissues is required.

## Background

There is now considerable evidence demonstrating that both pre- and postnatal exposure to beneficial microbial signals can heavily influence functional immune development during early life, resulting in the protection against future inflammatory disease^1-3^. The principal target of this beneficial immunostimulation appears to be the innate immune system^4, 5^, and the mechanisms driving protection are consistent with the newly emerging paradigm of “innate immune training”, whereby certain classes of microbial stimuli can alter the functional state of innate immune cells, leading to the optimisation of immunocompetence^6^. While immune training focusses on the phenotypic and transcriptional profiles of several prototypical innate populations^6^, further characterisation of the downstream adaptive response is critical in understanding the greater implications of innate immune training in the protection against future inflammatory disease susceptibility. As a result, the protective mechanisms remain incompletely understood. To address this requirement, we have developed and optimised a novel 19-parameter flow cytometry panel to comprehensively and simultaneously characterise distinct T-cell and B-cell subsets localised within lymphoid tissues of BALB/c neonatal mice following treatment with a microbial-derived immunomodulator.

The developmental phase of this flow cytometry panel involved the prioritisation of T-cell and B-cell subsets central to the maintenance of immunological homeostasis, as based on the current literature and forerunner studies. As such, a degree of emphasis was placed on effector, regulatory and memory subsets within T-cell and B-cell populations. In regards to T-cells, the conversion of peripheral naïve CD4^+^ T-cells to T-effector (Teff) cells is denoted by upregulation of the activation marker CD25, while concomitant upregulation of both CD25 and intracellular Foxp3 expression is essential for the peripheral induction of regulatory T cells (Treg)^7^, a process previously recognised in the protection against allergic airways inflammation following microbial-derived immunomodulation^8, 9^. Furthermore, the expression of CD44 on splenic Treg has been implicated in promoting enhanced function^10^. Following activation and contraction, CD4^+^ T-cells transition towards a memory phenotype via the gradual upregulation of CD44 expression in parallel with transient expression of CD62L, driving the establishment of a dynamic repository of central memory (T_CM_) and effector memory (T_EM_) T-cells^11, 12^. In addition to establishing peripheral memory, activated CD4^+^ T cells have the capacity to upregulate extracellular expression of CXCR5, inducible costimulator (ICOS) and programmed cell death protein 1 (PD-1)^13, 14^, resulting in the generation of a highly specialised population of T follicular helper (T_FH_) cells which accumulate within the B-cell follicles of secondary lymphoid organs and provide essential activation signals for the development of antigen-specific humoral immunity^15^. A separate subset of thymic-derived cells that share homology with the T_FH_ phenotype in addition to Foxp3 and bimodal CD25 expression, termed follicular regulatory T (T_FR_) cells, have also been identified, however this subset has been attributed to the inhibition of T_FH_ activity and subsequent generation of humoral immunity^16, 17^. The immunophenotypic characterisation of B-cell subsets for this OMIP was centred around the classic expression of CD19 and B220. To maximise the capacity of a 5-laser BD LSRFortessa™, CD19 (B-cell) and CD4 (T-cell) antibodies were conjugated to the same fluorochrome, since co-expression is absent in single cell analysis. Within secondary lymphoid tissues, the antigen-specific activation of B-cells involves the constitutive upregulation of major histocompatibility complex class-II (MHC-II; mouse I-A/I-E) and CD80 expression, in conjunction with the secretion of both immunoglobulin (Ig) M and IgD^18, 19^. Crucial to the application of this OMIP, previous studies have demonstrated that microbial-derived immunomodulation of BALB/c mice drives splenic B-cell activation and enhanced antibody production^20^. Following antigen-specific activation, B-cells upregulate Synd-1 expression and differentiate into the two major classes of antibody-secreting cells; the rapidly produced and short-lived plasmablasts (PB) and the short-lived peripheral plasma cells (PC), both of which have the capacity to secrete IgM^21, 22^. A major difference between these two antibody-secreting subsets however is the absence of classic mature B-cell markers CD19, B220 and MHC-II on plasma cells^22, 23^. The eventual transition of B-cells towards a memory phenotype results in the loss of Synd-1 expression with parallel upregulation of programmed cell death protein 1 ligand 2 (PD-L2), generating a long-lived secondary lymphoid population that can rapidly differentiate into antibody-secreting cells upon re-stimulation^24-26^.

Panel optimisation was performed on a BD LSRFortessa™, with all fluorochrome-conjugated antibodies (Table 2) titrated during the optimisation phase. Prior to multicolour extracellular staining, splenocytes were incubated in Fc Block™ (Purified recombinant CD16/32) to inhibit non-antigen-specific binding of fluorochrome-conjugated antibodies to the nonpolymorphic epitope of FcγIII (CD16) and FcγII (CD32) receptors expressed on multiple myeloid populations and B-cells. A representative gating strategy to delineate the T-cell and B-cell subsets described above is detailed in Figure 1. Briefly, splenocytes were first gated on side-scatter (SSC) and forward-scatter (FSC) parameters (Figure 1A) to remove sample debris, followed by single-cell gating (Figure 1B) to remove doublets. Gating was then performed on viable CD45^+^ cells (Figure 1C) to remove dead/dying cells and stromal cells from the analysis. The primary T-cell/B-cell separation involved delineation of TCRβ and CD4/CD19 expression (Figure 1D). Double positive cells were classified as CD4^+^ T-cells, as CD19^+^ B-cells will be present within the TCRβ^-^ population (Figure 1D). An additional TCRβ^-^CD4/CD19^-^ gate was included to enable the characterisation of B220^-^Synd-1^+^I-A/I-E^-^IgM^+^ PC (Figure 1E). CD19^+^ B-cells were then defined as B220^lo/+^Synd-1^+^I-A/I-E^+^IgM^+^ PB (Figure 1F), B220^+^Synd-1^-^CD80^+^PD-L2^-^I-A/I-E^+^IgM^+^IgD^+^ activated B-cells (Figure 1G) and B220^+^Synd-1^-^CD80^+^PD-L2^+^IgM^+^IgD^-^ memory B-cells (Figure 1H). CD4^+^ T-cells were defined as CD62L^+^CD44^-^ naïve T-cells (Figure 1I), CD62L^+^CD44^+^ T_CM_ (Figure 1I), CD62L^-^CD44^+^ T_EM_ (Figure 1I), CD25^+^Foxp3^-^ Teff (Figure 1J), CD25^+^Foxp3^+^ Treg (Figure 1J), CXCR5^+^ICOS^+^PD-1^+^ T_FH_ (Figure 1K) and CXCR5^+^ICOS^+^PD-1^+^CD25^+/-^Foxp3^+^ T_FR_ (Figure 1L).

**Table 1.** Summary Table

|  |  |
| --- | --- |
| Purpose | Comprehensive immunophenotyping of T-cell and B-cell subsets |
| Species | Mouse |
| Cell types | Lymphoid tissue single cell suspensions |
| Cross-reference | OMIP-031, OMIP-032, OMIP-054, OMIP-061 |

**Table 2.** Reagents used for OMIP

| Specificity | Fluorochrome | Clone | Purpose |
| --- | --- | --- | --- |
| PD-L2 (CD273) | BUV395 | TY25 | Memory B cells |
| IgD | BUV496 | 217-170 | Activated B cells |
| CD44 | BUV737 | IM7 | T cell subsets |
| ICOS (CD278) | BV421 | 7E.17G9 | T Follicular cells |
| PD-1 (CD279) | BV480 | J43 | T Follicular cells |
| FVS575 | BV605 | N/A | Viable cells |
| CD80 | BV650 | 16-10A1 | Activated B cells |
| IgM | BV711 | R6.60.2 | B cell subsets |
| CD4 | BV786 | RM4-5 | CD4<sup>+</sup> T cells |
| CD19 | BV786 | 1D3 | B cell subsets |
| Synd-1 (CD138) | BB515 | 145-2C11 | Plasmablasts/Plasma cells |
| TCRβ | BB700 | H57-597 | Pan-T cells |
| Foxp3 | PE | FJK-16s | Regulatory T cells |
| B220 (CD45R) | PE-CF594 | RA3-6B2 | B cell subsets |
| CD25 | PE-Cy5 | PC61 | Activated T cells |
| CXCR5 (CD185) | PE-Cy7 | 2G8 | T Follicular cells |
| I-A/I-E | AF647 | M5/114.15.2 | B cell subsets |
| CD62L | AF700 | MEL-14 | T cell subsets |
| CD45 | APC-H7 | 30-F11 | Pan-leukocyte |

**Figure 1.**
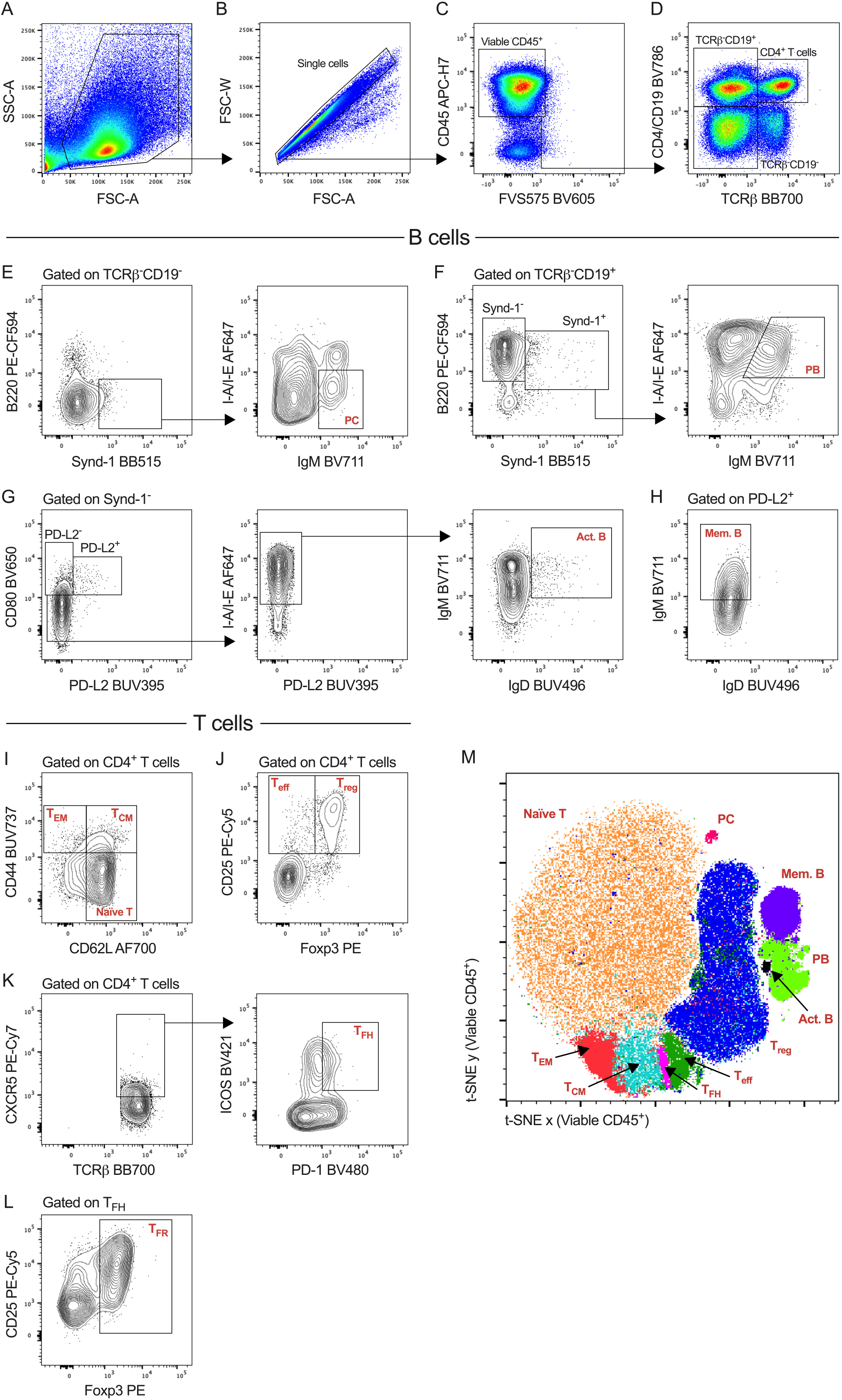
Overview of 19-parameter gating strategy developed for the characterisation of T-cell and B-cell subsets within freshly isolated splenocytes from 21-day-old neonatal BALB/c mice. 1×10^6^ splenocytes were incubated in FC Block™, followed by fixable viability stain (FVS) and a 17-parameter extracellular antibody cocktail containing 10% Brilliant Stain Buffer Plus (BD Biosciences). Intracellular staining was performed following fixation-permeabilisation of extracellular stained splenocytes. Data was acquired on a BD LSRFortessa™ (BD Biosciences). **(A-C)** Removal of cellular debris, doublets, non-viable cells and stromal cells. **(D)** Primary delineation of TCRβ^-^CD19^+^, TCRβ^+^CD4^+^ and TCRβ^-^CD4/CD19^-^ cells. **(E-L)** Characterisation of **(E)** plasma cells, **(F)** plasmablasts, **(G)** activated B-cells, **(H)** memory B-cells, **(I)** naïve, effector memory and central memory T-cells, **(J)** effector and regulatory T-cells, **(K)** T follicular helper cells and **(L)** follicular regulatory T-cells. All plots are representative of individual samples. **(M)** Algorithm-based t-SNE analysis of viable CD45^+^ splenocytes with manual gating overlayed to distinguish T-cell and B-cell subset clusters. t-SNE plot was generated via the concatenation of 300,000 total viable CD45^+^ cells from 12 individual samples with the following parameters; iterations: 5000, perplexity: 30, learning rate (eta): 21102.

Manual gating was validated by t-Distributed Stochastic Neighbor Embedding (t-SNE) of viable CD45^+^ splenocytes (gated in Figure 1C) to construct algorithm-based subset clusters for the visualisation of surface and intracellular receptor co-expression profiles of T-cell and B-cell subsets (Figure 1M). As expected, T-cell and B-cell subpopulations are restricted within distinct clusters, however PC are spatially isolated from memory B-cells, activated B-cells and PB due to their lack of CD19 expression (Figure 1D and E).

### Similarities to other OMIPs

The OMIP described here shares a small degree of marker similarity (TCRβ, CD4, CD44, CD62L, PD-1, CD19, B220) with OMIP-031^27^, OMIP-032^28^ and OMIP-061^29^, which are focussed on immunologic checkpoint expression on murine T-cell subsets, the characterisation of innate and adaptive populations within the murine mammary gland and murine antigen-presenting cells, respectively. While both OMIP-031 and OMIP-032 characterise TCRβ^+^CD4^+^ effector and memory T-cell subsets based on a combination of CD44 and/or CD62L expression, OMIP-032 employs an additional CD19^+^ gate to delineate B-cells. OMIP-061 utilised B220 to identify B-cells. A distinct difference between these OMIPs and the OMIP described here is that our panel was developed for the sole purpose of comprehensively immunophenotyping T-cell and B-cell subsets in lymphoid tissues, and we therefore include an additional 12 markers to allow the simultaneous characterisation of 4 major B-cell and 7 major T-cell populations within a single sample. The OMIP described here also exhibits minor overlap with OMIP-054^30^, however our panel was developed to maximise the potential of a 5-laser BD LSRFortessa™ in facilities without the capacity to perform mass cytometry.

## Acknowledgements

The authors would like to thank Steven Roberts and Dr Andrew Lim of BD Biosciences (Australia) for their valuable advice during the initial design of this panel.

